# LncRNA H19X is required for placenta development and angiogenesis through regulating a noncoding RNA regulatory network

**DOI:** 10.1101/2020.11.19.389940

**Authors:** Li Tongtong, Yacong Cao, Yanting Zou, Ye Yang, Wang Ke, Huang Gelin, Li Xiaoliang, Zheng Rui, Tang Li, Lv Jiao, Yang Ming, He Jiabei, Zhang Xiaohu, Bai Shujun, Li Qintong, Qin Lang, Zhao Xiao Miao, Xu Wenming

## Abstract

H19X is a lncRNA specifically expressed in the placenta, whose expression is induced by hypoxia. H19X overexpression promoted trophoblast proliferation and invasion, while its knockdown or knockout inhibited trophoblast proliferation and invasion. Mechanistically, we demonstrated that reciprocal regulation exists with miR-424/miR-503 in the control of genes related to placental development and angiogenesis, including VEGF and VEGFR2. H19X inhibited ubiquitination of PIWIL1, thereby maintaining its stability and homeostatic expression of piRNAs. PIWIL1 overexpression rescued the defects of cell behavior caused by H19X KO. H19X deletion led to compromised HIF-1A/HIF-2A expression, which was correlated with the dysregulation of downstream genes under hypoxic conditions. CRISPR/Cas-9 knockout of H19X in animals led to defective placenta differentiation and compromised embryo development under hypoxic conditions. Western blotting showed reduced expression levels of PIWIL1 as well as angiogenesis marker genes, including VEGF and VEGFR2, in H19X KO mice. Thus, this study provides evidence of an unexpected link among lncRNA, miRNA, PIWIL1-related piRNA, and angiogenesis in placentation, the dysregulation of which leads to poor placental development and embryo loss under hypoxic conditions.

## Introduction

Long non-coding RNAs (lncRNAs) are a group of RNA molecules longer than 200 bp that do not encode proteins (Necsulea *et al*, 2014). Emerging evidence indicates that lncRNAs play important roles in embryo development, cell cycle regulation, and tumor development. Recently, it has also been shown that lncRNAs are critically involved in placental physiology, including X chromosome inactivation, imprinting, and DNA methylation (Hemberger *et al*, 2020; Penkala *et al*, 2016). The diverse involvement of lncRNAs in different cellular functions is in accordance with their tissue-specific expression patterns (Ang *et al*, 2020; Batista & Chang, 2013). For example, it has been shown that an lncRNA highly expressed in placenta, H19, plays versatile roles in placental biology and embryogenesis (Troy *et al*, 2012). Interestingly, a disease association analysis identified an lncRNA, HELLP, as being linked to the pathogenesis of HELLP syndrome, a disease occurring during pregnancy caused by an abnormality of the third trimester fetal placenta (van Dijk *et al*, 2012). These results and others indicate that lncRNAs play a critical role in placental pathophysiology.

Compared with other non-coding RNA molecules, such as miRNAs, lncRNAs have several distinct characteristics. First, lncRNAs show more tissue-specific expression patterns. Second, lncRNAs have more complex regulatory mechanisms; for example, lncRNAs can act as decoys by binding with proteins to form complexes. Third, lncRNAs can form complexes with proteins as scaffolds to direct other proteins to exert their regulatory functions. Fourth, lncRNAs have the ability to form RNA-RNA double helices, and therefore affect the functions of other non-coding RNAs (Robinson *et al*, 2020). It also became obvious that some lncRNA could encode short peptides, and therefore contribute to the regulation of down-stream pathway (Huang *et al*, 2017). Although their regulation is complex, it is widely accepted that lncRNA-protein complexes play important regulatory roles in downstream gene expression. One well-known lncRNA involved in placental development is H19, which is an imprinted and maternally expressed lncRNA important for embryo development (Keniry *et al*, 2012; Smits *et al*, 2008). Interestingly, H19 contains miR-675 in its first exon, and therefore, functions as a miRNA sponge to regulate IGF-1 to direct embryo development. H19 is also important for trophoblast migration as it targets TβR3 (Zuckerwise *et al*, 2016). Therefore, it is well accepted that lncRNAs play multiple roles during placentation.

In a recent study combining high-throughput sequencing with bioinformatics analysis, H19X was identified as a placenta-specific lncRNA on the X chromosome, with an expression mode that is conserved from mice to monkeys and humans (Necsulea *et al.*, 2014). Other transcriptome analysis indicated that H19X is specifically expressed in the placenta, where its expression is co-regulated with the miR-15/16 family (Wang *et al*, 2019b). Specifically, the first exon and first intron region of H19X were found to overlap with miR-424 and miR-503, which belong to the miR-15/16 family, play important roles in angiogenesis, and have been implicated in the regulation of trophoblast behavior. In our previous study, we showed that miR-15b is abundantly expressed in trophoblasts and inhibits the invasion of trophoblasts by regulating Ago2 expression (Yang *et al*, 2016). In another recent study, we showed that Dicer1 and miRNAs in the placenta are highly expressed in trophoblasts, and both were secreted from trophoblasts, regulated endothelial cell behavior through extracellular vesicles, and modulated angiogenesis during placentation (Muys *et al*, 2016; Tang *et al*, 2020). However, it is not known whether H19X is involved in placentation and embryo development *in vivo*. The present study was conducted to investigate whether H19X plays a role in placental regulation and to explore the underlying mechanism. Our results showed that H19X plays an important role in trophoblast behavior, and we identified major defects resulting from *in vivo* deletion of H19X. Furthermore, novel functions, including maintaining PIWI stability and reciprocal regulation with HIF-2A, revealed the critical role of H19X in mediating placental development and angiogenesis, demonstrating that H19X is critical for linking miRNA and piRNA expression during placentation.

## Results

### H19X is expressed specifically in the placenta of humans and mice

As H19X is a newly identified lncRNA, we first confirmed its expression pattern in different mouse tissues by *in situ* hybridization (ISH) and real-time PCR (RT-PCR). A previous study found H19X expression shifts from the testis to the placenta during the appearance of placenta in animal evolution (Necsulea *et al.*, 2014), while anothers showed that H19X is highly expressed in trophoblast cells (Wang *et al.*, 2019b). Fluorescence *in situ* hybridization (FISH) was used to evaluate the localization of H19X in the placenta and trophoblast cells. The results confirmed that H19X was localized to the villus of the human placenta (Figure 1A). Using ISH, H19X was found to be mostly expressed in the cytoplasm of HTR-8/SVneo cells (Figure 1B). Furthermore, RNA profiling of mouse tissues showed that H19X is highly and specifically expressed in the placenta compared with other tissues, including the testes and brain (Figure 1C). These results suggest that H19X could potentially play a critical role in placental development.

**Figure 1.**
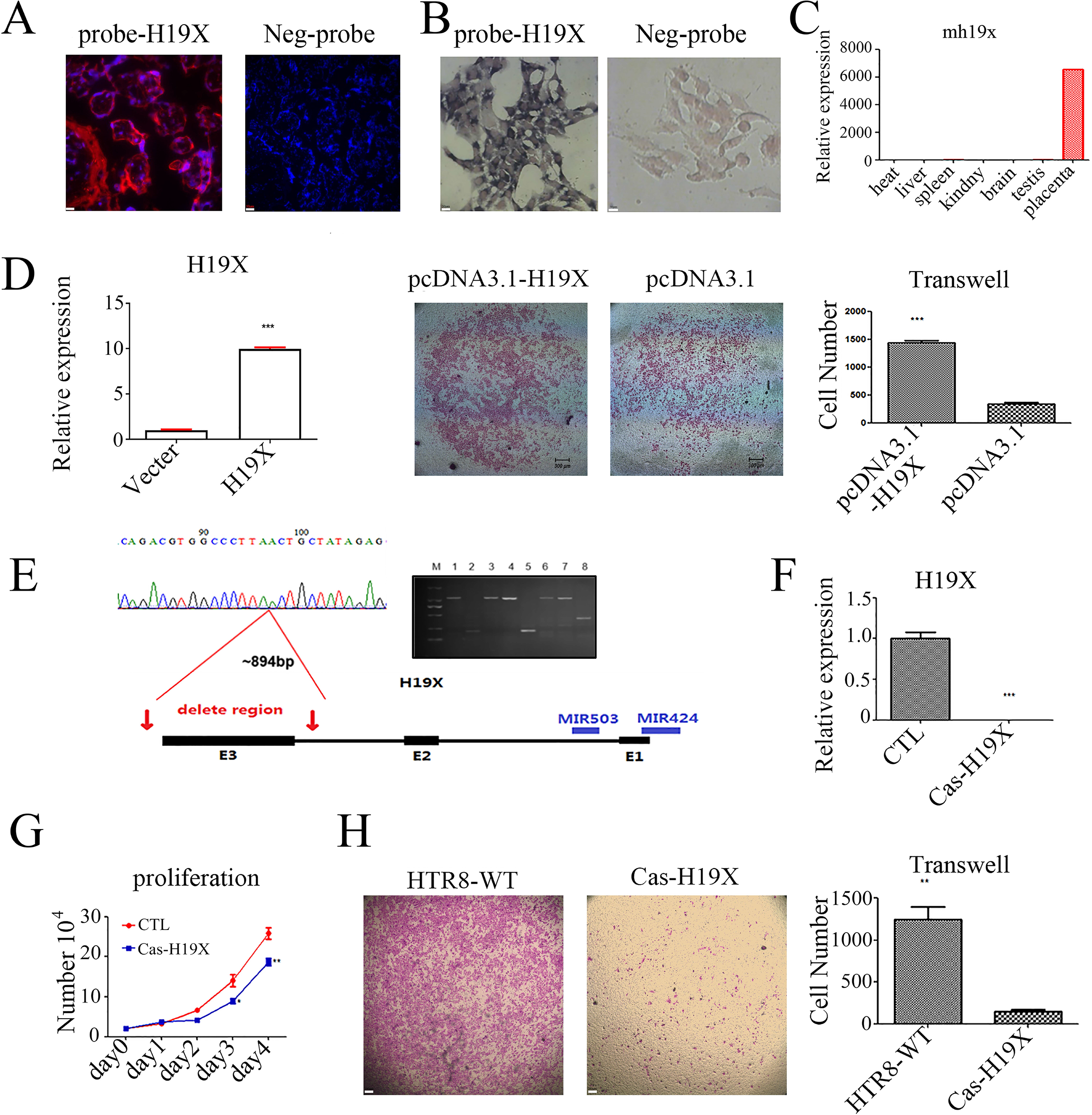
H19X is highly expressed in the trophoblasts of the placenta and is required for normal trophoblast behavior. (A) ISH staining showed that H19X is highly expressed in the cytoplasm of extravillous trophoblast cells. The right panel is an image with the negative control probe. (B) ISH staining of HTR-8SV/neo cells confirmed cytoplasmic staining of H19X. The right panel shows the negative control. (C) Real-time PCR of multiple mouse tissues showed that H19X is specifically expressed in the placenta. (D) The left panel shows that H19X is significantly increased in the H19X overexpression group, and the right panel shows that H19X overexpression led to increased trophoblast invasion. (E) Deletion of exon 3 of H19X. The right gel shows a positive clone from the CRISPR/Cas-9 knockout. (F) Real-time PCR showing that H19X expression was abolished in the knockout cell line. ***p < 0.001. (G) Cell proliferation curve showing that deletion of H19X reduced cell proliferation. (H) Matrigel invasion assay showing that CRISPR/Cas-9 deletion of H19X resulted in significantly reduced cell invasion. The right panel shows the statistical results. *p < 0.05, **p < 0.01.

### H19X plays important roles in human trophoblast cell proliferation and invasion

Since H19X is highly expressed in trophoblast cells, we investigated whether H19X affects trophoblast cell function. We measured the proliferation, invasion, and angiogenesis ability of HTR-8/SVneo cells after manipulating the expression of H19X. Overexpression of H19X was confirmed by real-time detection of H19X transcripts after transfection of H19X expression plasmids, which showed a significant increase in expression (Figure 1D). H19X overexpression was found to induce trophoblast invasion in Matrigel (Figure 1D). We also established an H9X knockdown model, and the whole transcriptome and small-interfering RNA sequence flowchart are shown in Supplementary Figure 1. When H19X was knocked down using shRNA, HTR-8/SVneo cells showed reduced cell invasion capability (Supplementary Figure 2). CRISPR/Cas-9 knockout is being increasingly used to validate the function of genes due to its simplicity and high efficiency compared with other techniques, such as RNAi^15^. Therefore, we used the CRISPR/Cas9 technique to knockout H19X in HTR-8/SVneo cells to further evaluate its function. Because a cluster of miRNAs, including miR424 and miR503 are located near the first exon in the 5′-terminus of H19X, and the third exon of human H19X is evolutionarily conserved, we decided to only knockout exon 3 of H19X in HTR-8/Svneo cells (Figure 1E). Electrophoresis indicated successful deletion in a single cell-derived colony, which was further confirmed by sequencing and RT-PCR analysis (Figure 1E, F). Consistent with the behavior of H19X knockdown cells, cell proliferation and invasion were largely inhibited in the Cas-H19X HTR-8/SVneo cells when compared to the corresponding behaviors of WT HTR-8/SVneo cells from the same passage (Figure 1G, H). These results indicate that H19X plays a critical role in regulating human trophoblast proliferation and cell invasion.

### H19X maintains miR-424/503 expression and the reciprocal regulation between H19X and miR-424

Since the miRNA cluster miR424/503 is located near the first exon of H19X (Figure 1E), we explored whether H19X affected the expression of the miR-424/503 cluster. H19X overexpression induced the expression of miR-503 and miR-424 (Figure 2A). Deletion of H19X with CRISPR/Cas-9 almost completely abolished the expression of H19X, while the expression of miR-503 and miR-424 was significantly reduced (Figure 2B). We evaluated the expression levels of pre-miR424 and pre-miR503 in Cas-H19X cells and found that their expression levels were also reduced (by approximately 60%), similar to the change in mature miR424 and miR503 expression levels following H19X knockout (Figure 2C). These results indicate that H19X may regulate miR424/503 at the transcriptional level. Conversely, to determine the potential roles of miR424/503 in H19X expression, we examined the expression level of H19X in HTR-8/SVneo cells after transfection of an miR424 mimic or miR503 mimic. Interestingly, we found that only miR424 affected H19X expression in a dose-dependent manner, while the miR503 mimic had no effect on H19X expression (Figure 2D). This result confirms the reciprocal regulation between H19X and miR-424 at the transcriptional level. To further determine whether knockdown of H19X affects the expression of miR-424/miR-503 and downstream targets, we used small RNA-seq and RNA-seq analysis of three independent lines of H19X knockdown cells (Supplementary Table 2 and 3). Expression levels were calculated, and the differentially expressed genes, miRNAs, and piRNAs between samples were identified (Supplementary Figure 3). Our small RNA sequencing results showed that knockdown of H19X inhibited the expression of miR-424 and miR-503 (Figure 2E). Analysis of the miRNA-target regulatory network showed that knockdown of H19X had a more significant impact on the target genes of miR-424-5p than on the target gene of miR-424* (Supplementary Figure 4), with similar results observed for miR-503 (Supplementary Figure 5). GO analysis further showed that the target genes of both miR-424 and miR-503 were related to various aspects of placentation and organ development (Supplementary Figures 4 and 5). miRNA-target analysis showed that miR-424 both upregulates and downregulates targets (Figure 2F). Analysis of the top 20 most significantly differentially expressed miRNAs showed that the target genes are involved in various aspects of organ development, including “vascular process in circulatory system”, “urogenital system development”, “stem cell differentiation”, et,al (Supplementary Figure 6). Taken together, these results indicate that deletion of H19X resulted in the downregulation of miR-424, which contributes to the dysregulation of target genes related to placental function and embryo development.

**Figure 2.**
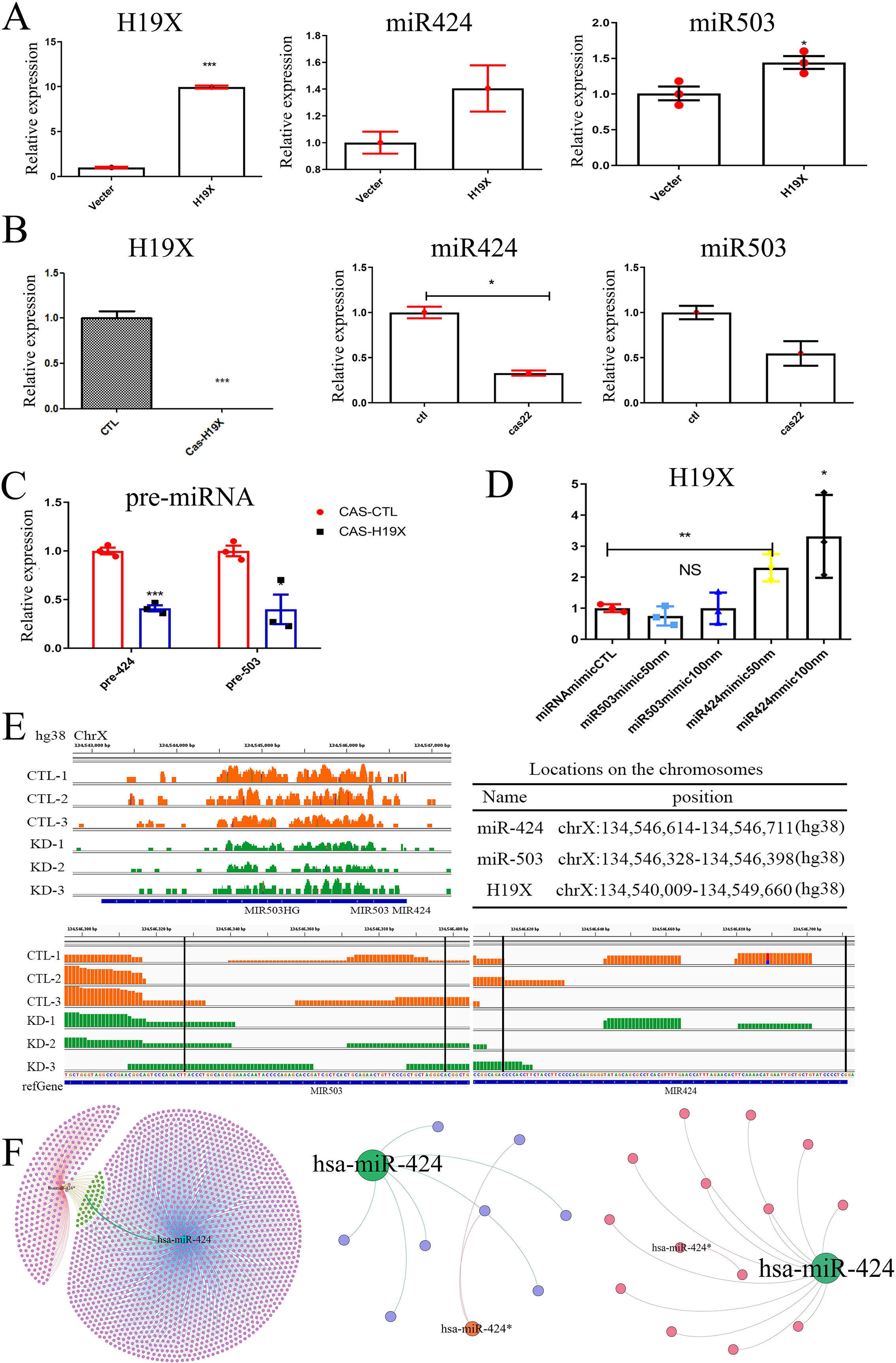
H19X expression and miR-424 expression are reciprocally regulated in trophoblast cells. (A) H19X upregulation in H19X-overexpressing cells. The expression levels of both miR-424 and miR-503 were also significantly increased. (B) H19X knockout in an HTR-8 cell line results in significantly decreased expression of miR-424 and miR-503. (C) Real-time PCR shows that deletion of H19X results in significant downregulation of pre-miR-424 and pre-miR-503 expression. (D) Real-time PCR shows that H19X expression is dose-dependently upregulated by miR-424 transfection but not by miR-503 transfection in HTR-8 cells. (E) The small RNA sequencing results showed that deletion of H19X results reduced miR-424 and miR-503 expression. (F) The miRNA-target regulatory network analysis showed that miR-424 regulated more targets. On the left, the upregulated DEGs are shown, while on the right, the downregulated DEGs are shown. hsa-miR-424 (mature sequence hsa-miR-424-5p): cagcagcaauucauguuuuga; hsa-miR-424* (mature sequence hsa-miR-424-3p): caaaacgugaggcgcugcua. *p < 0.05; **p < 0.01; ***p < 0.001.

### H19X interacts with PIWIL1 and regulates PIWIL1 protein levels

Since H19X was found to play important roles in trophoblast behavior, and most lncRNAs exert their functions via protein binding, we used an RNA pull-down assay to identify proteins that interact with H19X. One significant protein band appeared exclusively in the H19X sample that was not present in the control beads only sample. Mass spectrometry was performed to determine the identity of the protein, which indicated that the protein was PIWIL1, a potential protein interacting with H19X (Figure 3A and Supplementary Table 4). Western blotting further confirmed that PIWIL1 interacts with H19X (Figure 3B). We then conducted an RNA immunoprecipitation (RIP) experiment to determine whether PIWIL1 could interact with endogenous H19X in HTR-8/SVneo cells. The retrotransposon Line-1 was used as a positive control due to the participation of PIWIL1 in piRNA-mediated retrotransposon silencing. The results confirmed that PIWIL1 interacted with H19X not only *in vitro* but also within cells (Figure 3C). To determine whether these interactions affect the levels of PIWIL1 protein, we evaluated PIWIL1 protein expression in H19X knockdown and knockout cells. Accordingly, a significant decrease in PIWIL1 expression was observed (Figure 3D). These results indicate that H19X interacts with PIWIL1 protein and maintains PIWIL1 protein expression in HTR-8/SVneo cells.

**Figure 3.**
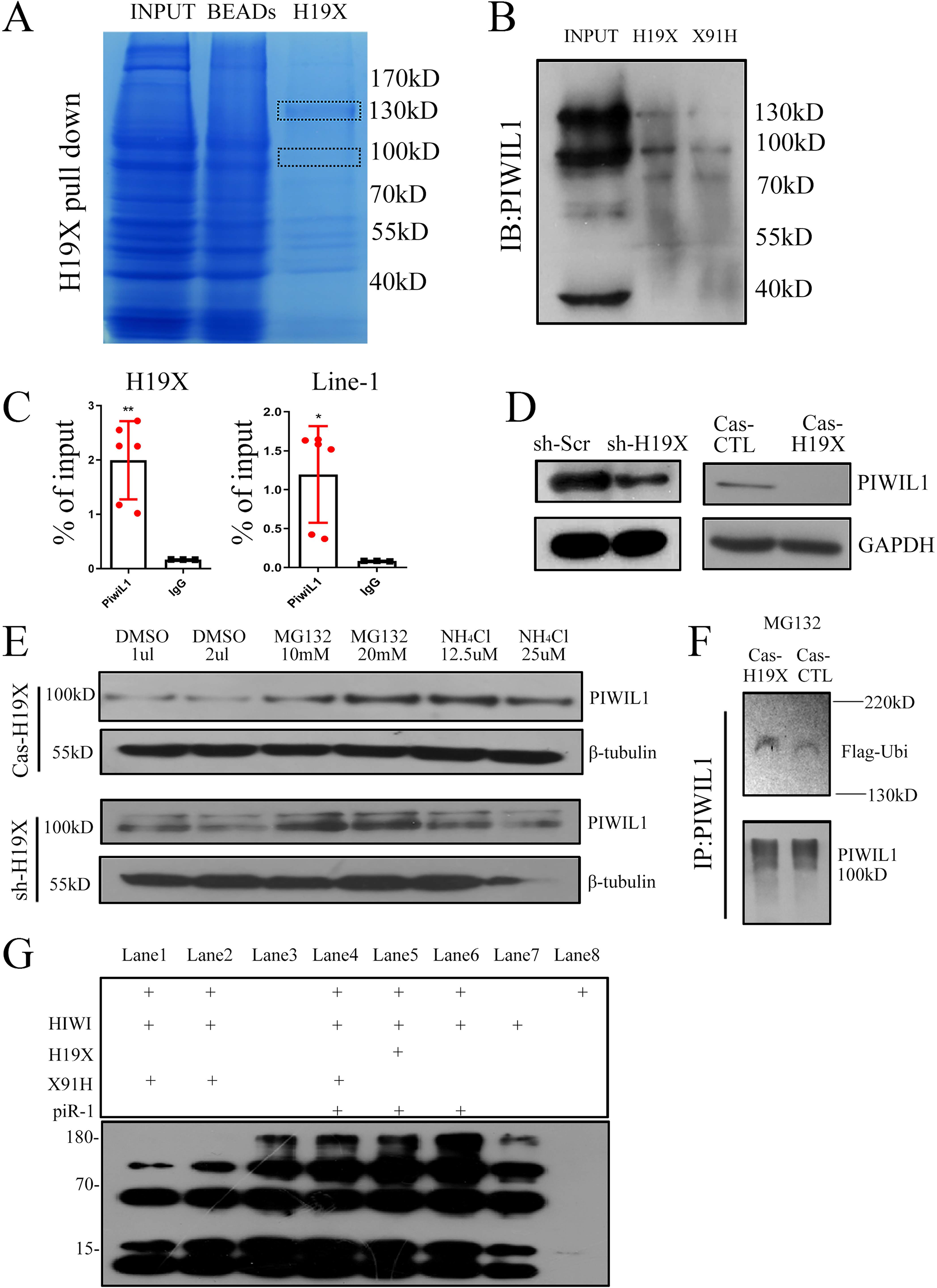
H19X binds directly to PIWIL1 and protects it from degradation by inhibiting its ubiquitination. (A) The results of the H19X pull-down assay show that H19X can pull-down a protein from cell pellets. Mass spectrometry identified the band around 100 kDa as PIWIL1, indicating that PIWIL1 is a possible target protein. (B) The results of western blotting using a H19X-labelled probe show that H19X can bind to PIWIL1. Anti-sense probe was used as the negative control in the binding assay. (C) RIP assay showing that Piwil1 immunoprecipitated H19X. Right panel shows the positive control, Line-1. *p < 0.05; **p < 0.01. (D) Silencing of H19X leads to reduced PIWIL1 expression, and CRISPR/Cas-9 KO results in diminished PIWIL1 expression. (E) Inhibition of the proteasome with MG-132 in H19X KO leads to increased PIWIL1 stability. (F) A ubiquitination assay shows that deletion of H19X leads to increased ubiquitination, as shown by probing with Flag-ubi and PIWIL1 protein, indicating that PIWIL1 ubiquitination is increased in H19X KO cells. (G) *In vitro* ubiquitination assay showing that adding piRNA-1 resulted in higher PIWIL1 protein expression.

Next, we investigated the mechanism by which H19X affects PIWIL1 expression. A wide range of mechanisms are involved in protein degradation, among which the ubiquitin-proteasome- and lysosome-mediated degradation play key roles. Therefore, we examined PIWIL1 protein levels after treating H19X-knockdown or -knockout HTR-8/SVneo cells with the proteasome inhibitor MG132 or the lysosome inhibitor NH_4_Cl. The results showed that 10–20 μM MG132 significantly enhanced PIWIL1 protein levels in both H19X knockdown and knockout cells (Figure 3E), indicating that H19X regulates PIWIL1 protein by inhibiting proteasome-mediated degradation, in accordance with previous studies showing that degradation of mouse testis PIWIL1 is mediated by the ubiquitin-proteasome pathway (Zhao *et al*, 2013). Then, we transfected 4×Flag-Ubiquitin into Cas-H19X and Cas-CTL cells and treated them with 20 μM MG132 to inhibit proteasome function, which led to the same PIWIL1 protein levels in both Cas-H19X cells and Cas-CTL cells, whereas ubiquitinated PIWIL1 levels were significantly increased in Cas-H19X cells (Figure 3F). These results confirm that H19X maintains PIWIL1 stability by inhibiting its ubiquitination. In previous studies, *in vitro* ubiquitination analysis indicated that the process of ubiquitination requires piRNAs (Gou *et al*, 2017; Zhao *et al.*, 2013). Adding piR-1 to the *in vitro* system further inhibited the ubiquitination process, as shown in Figure 3G. Taken together, these results show that H19X maintains the stability of PIWIL1, which plays a critical role in piRNA production during placental development.

### PIWIL1 plays important roles in trophoblast behavior

PIWIL1 is a germ cell-expressed gene that is mainly involved in germ cell development, but has never been shown to be expressed in placental tissue. Thus, we first checked whether PIWIL1 was expressed in placental tissue and trophoblast cells. Using immunohistochemistry, we confirmed that PIWIL1 is highly expressed in human testes and the human villus of the placenta (Figure 4A). The major PIWIL1 homologues in mice are MIWI1 and MILI(Fenner, 2017). In the present study, RNA analysis showed that MIWI is more dynamically regulated, and the highest expression was detected on E13.5 day (Figure 4B). To investigate whether PIWIL1 plays a role in trophoblast behavior, we designed a lentivirus to specifically knockdown PIWIL1 in HTR-8/SVneo cells and confirmed the knockdown effect by western blotting (Figure 4C). Knockdown of PIWIL1 was found to reduce cell invasion, based on the results of an invasion assay (Figure 4D). Similarly, the results of a cell proliferation assay showed that cell proliferation was significantly reduced after H19X knockdown (Figure 4E). To further confirm that PIWIL1 is downstream of H19X regulation, we expressed PIWIL1 in H19X Cas-9 deletion cells, and both the cell proliferation and invasion assays indicated that PIWIL1 rescued the cell proliferation and invasion defects caused by H19X deletion (Figure 4F,G), confirming that the function of PIWIL1 is downstream of H19X, at least partially, to mediate the function of H19X in placental development.

**Figure 4.**
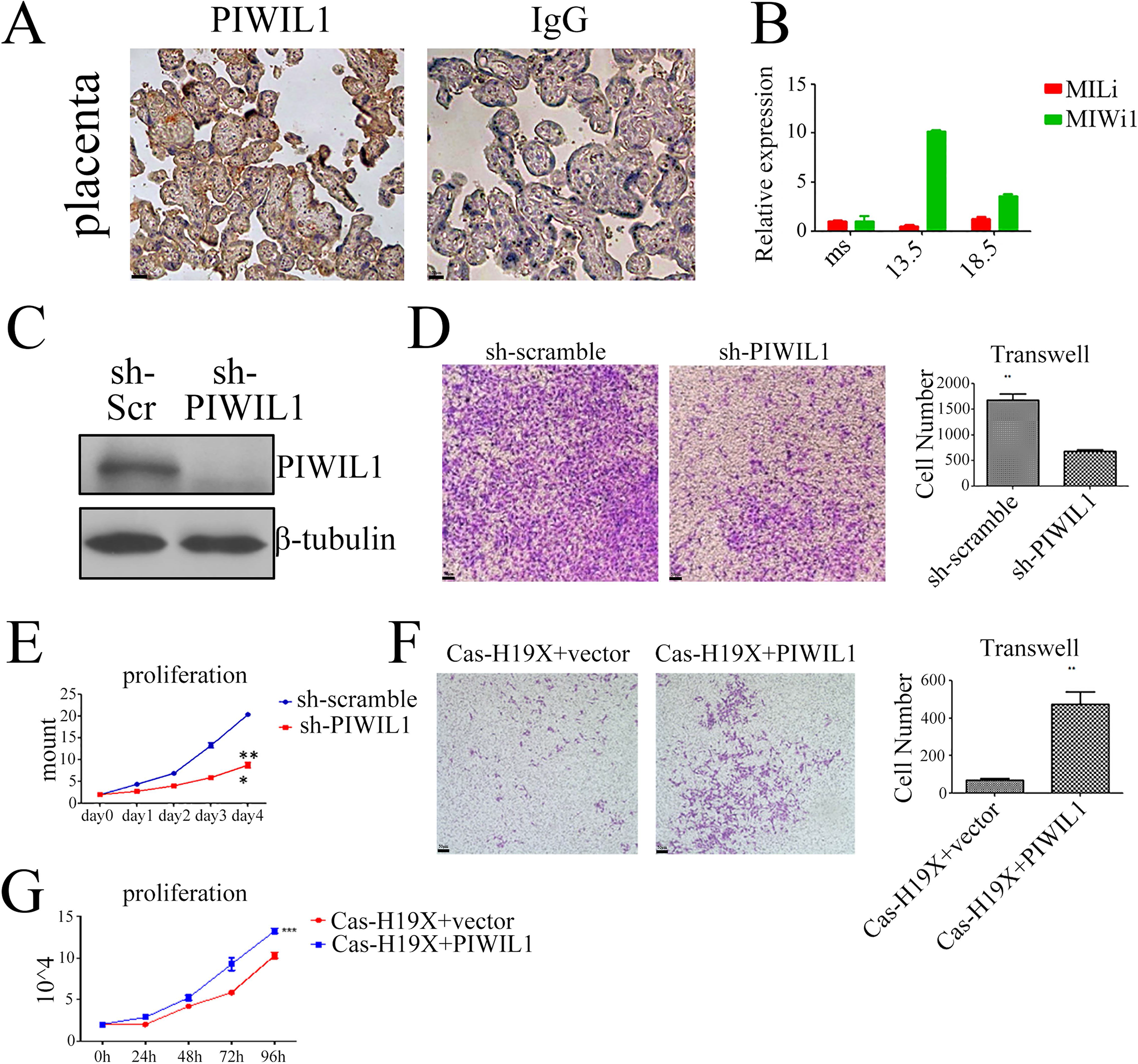
PIWIL is essential for the invasion and proliferation of trophoblasts. (A) IHC staining shows that PIWIL1 is expressed in trophoblasts (IgG, negative control). (B) The mouse homologue of PIWIL1, miwi1, is dynamically expressed during placental development. Ms, E6 mouse embryo. Miwi1 shows the highest expression levels in embryo on E13.5. Miwi1 expression levels begin to decrease on E18.5. (C) Western blot showing that PIWIL1 KO results in diminished PIWIL1 expression. (D) Cell invasion assay showing that PIWIL1 KO leads to a reduced number of migrated cells in the invasion assay. (E) Cell proliferation assay showing that PIWIL1 KO reduces cell proliferation. (F) Transfection of PIWIL1 results in increased cell invasion, as shown by an increased number of invaded cells in the PIWIL1 overexpression group compared with the KO control. (G) Transfection of PIWIL1 in H19X KO cells results in increased cell proliferation when compared with the KO control.

### H19X maintains the immune suppressive status of trophoblasts via piRNA-mediated regulation

The major function of PIWIL1 is to regulate the production of piRNAs. Thus, it is involved in piRNA-mediated regulation of downstream targets. To further determine whether H19X could affect piRNA production, and therefore downstream functions, we used small RNA sequencing to evaluate piRNA expression in a H19X KD cell model. The results showed that when H19X KD HTR-8 cells were compared with a control cell line, 220 piRNAs were differentially expressed; 88 piRNAs were upregulated and 132 piRNA were downregulated (Table S3). The most significantly expressed piRNAs were analyzed, and the 20 most significantly changed piRNAs and the piR-target regulatory networks of upregulated and downregulated piRNAs are shown in Figure 5A and B. Using GO analysis, immune activation and related functions were found to be enriched among both the upregulated and downregulated target genes (Figure 5C, D). Taken together, these results show that H19X inhibits PIWIL1 ubiquitination and therefore affects piRNA production in trophoblast cells, and may modulate the immune response of trophoblasts. There are many genes with overlapping piRNAs. Functional enrichment analysis showed that a significant number of genes were related to RNA metabolism (Supplementary Figure 7). GOanalysis of the target genes of the differentially expressed piRNAs showed that the major function is in agreement with the essential role of the placenta in supporting embryo development (Supplementary Figure 8). Overall, these results show that H19X affects piRNA production by regulating PIWIL1 stability. Thus, piRNAs may be important for the phenotype observed in the H19X KO cell model.

**Figure 5.**
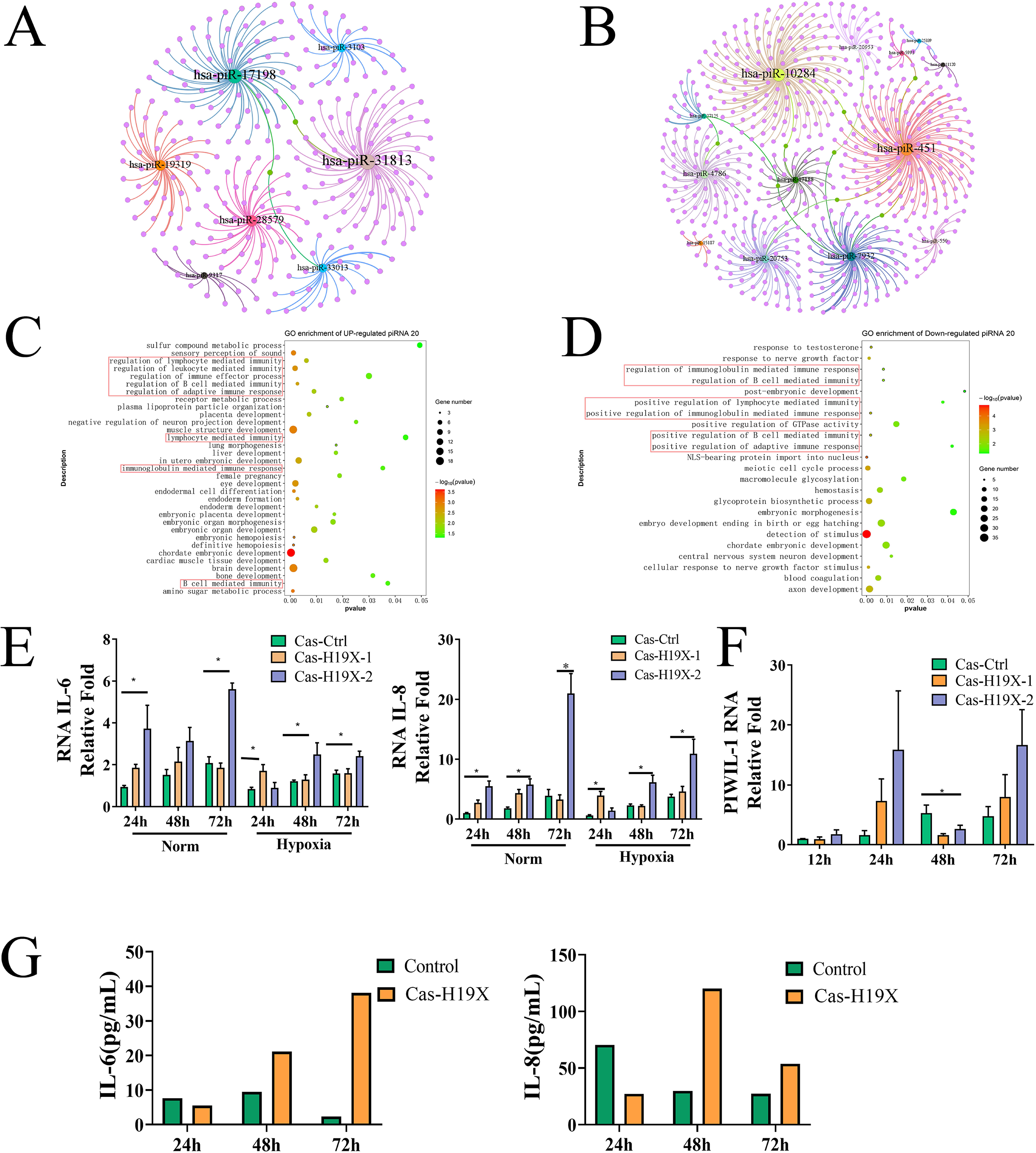
Small RNA sequencing shows that dysregulated piRNA expression is related to immune activation in trophoblast cells. (A, B) The top 20 most significantly differentially expressed piRNAs and their target genes. Left, upregulated piRNAs; right, downregulated piRNAs. (C, D) GO analysis of the target genes of the differentially expressed piRNAs shows that both the upregulated and downregulated piRNAs regulate genes involved in immune-related pathways. (E) Real-time PCR shows that CRISPR/Cas-9 deletion of H19X results in increased IL-6 and IL-8 expression and downregulated PIWIL1 expression. (G) Cytokine array show that CRISPR/Cas-9 KO of H19X leads to increased secretion of IL-6 and IL-8 into HTR-8 cell conditional medium. Cas-H19X-1 and Cas-H19X-2 are two independent CRISPR/Cas-9 KO cell clones.

Since the immunosuppressive status of the placenta is important for trophoblast behavior during trophoblast development, we evaluated cytokine and PIWIL1 expression under normal and hypoxic conditions. IL-6 and IL-8 are major cytokines involved in immune regulation in the placenta. Our results indicated that knockout of H19X induced the mRNA expression of IL-6/IL-8. PIWIL1 expression was increased after 24 h, but was then reduced after 48 h (Figure 5E). Using a cytokine array, we showed that after deletion of H19X, the secretion of IL-6 and IL-8 into trophoblast cell medium was significantly induced (Figure 5F). These results indicate that H19X deletion led to increased IL-6/IL-8 production and secretion, which may result in an overactivated immune response following knockdown of H19X in trophoblast cells.

Since miRNAs and piRNAs are small RNAs that execute their function by regulating target gene expression, we conducted transcriptome sequencing and evaluated the changes in the transcriptome after knockdown of H19X. GO analysis of the transcriptome is shown in Supplementary Figure 9. Interestingly, the enriched biological functions among the downregulated genes included “response to oxygen” and “positive regulation of sprouting angiogenesis” (Supplementary Figure 9). To examine the extent to which piRNAs and miRNAs are involved in differentially expressed gene (DEG) expression, the target genes of the piRNAs, miRNAs, and DEGs were analyzed. The analysis showed that 48.5% of the target genes of the piRNAs were downregulated (Supplementary Figure 10). Furthermore, more miRNA target genes (84.3%) were downregulated, and the GO analysis indicated that the function of these genes is related to “response to oxygen levels” and related pathways (Supplementary Figure 11). The overlap between the target genes of the differentially expressed miRNAs and DEGs is shown in Supplementary Figure 11. Interestingly, the target genes of both the miRNAs and piRNAs were analyzed, and the overlapping genes were found to be broadly involved in embryo development, with “VEGF production” and “response to oxygen levels” among the top downregulated biological functions (Supplementary Figure 12).

### H19X expression is regulated by HIF-1A/HIF-2A under hypoxic conditions

Previous studies have shown that H19X is regulated by hypoxia. Moreover, it is well known that HIF1a/HIF2A is a master transcriptional factor involved in the regulation of trophoblast behavior under hypoxic conditions. It has also been shown that some miR-15/miR16 molecules, including miR-107, can regulate the HIF pathway. As such, we evaluated whether deletion of H19X affects the expression of HIF-A under hypoxic conditions. and whether hypoxia leads to changes in HIF factors. An increase in H19X expression was observed after 24 and 48 h of hypoxia treatment, which was decreased after 72 h (Figure 6A). Hypoxia treatment also led to the downregulation of PIWIL1 protein expression (Figure 6B). We then investigated whether knockout of H19X affected trophoblast cell invasion under hypoxic conditions, and found that H19X knockout also compromised cell invasion (Figure 6C). To further investigate whether H19X affects HIF-1a/HIF-2a-regulated pathways, we evaluated RNA transcriptome data from the H19X knockdown cell line and analyzed the correlation of HIF-1A and HIF-2A expression levels with gene expression in H19X knockdown cells. The correlation analysis indicated that both HIF1A and HIF-2A were correlated with the upregulated and downregulated genes (Figure 6D). Interestingly, the correlation analysis showed that, compared with HIF-1A, HIF-2A was more significantly correlated with both the upregulated and downregulated genes. This analysis also indicated that HIF-2A may be involved in the transcriptional regulation of H19X (Figure 6E). Then, we checked the transcriptome results and confirmed that HIF-2A targets genes in the TRRUST database. Our results indicated that HIF-2A could regulate a significant number of the genes regulated by H19X. One of the most significant pathways is the VEGF-VEGFR2 pathway (Figure 6F, G). A conserved motif in the HIF-2A binding sites was determined (Figure 6H). Then, the H19X promoter sequence was extracted from UCSC to identify the transcription factors that bind to this region. The results showed that the HIF2A motifs exist in the H19X promoter. Taken together, these transcriptome data indicate that HIF1A/HIF-2A regulation plays a critical role in mediating H19X function.

**Figure 6.**
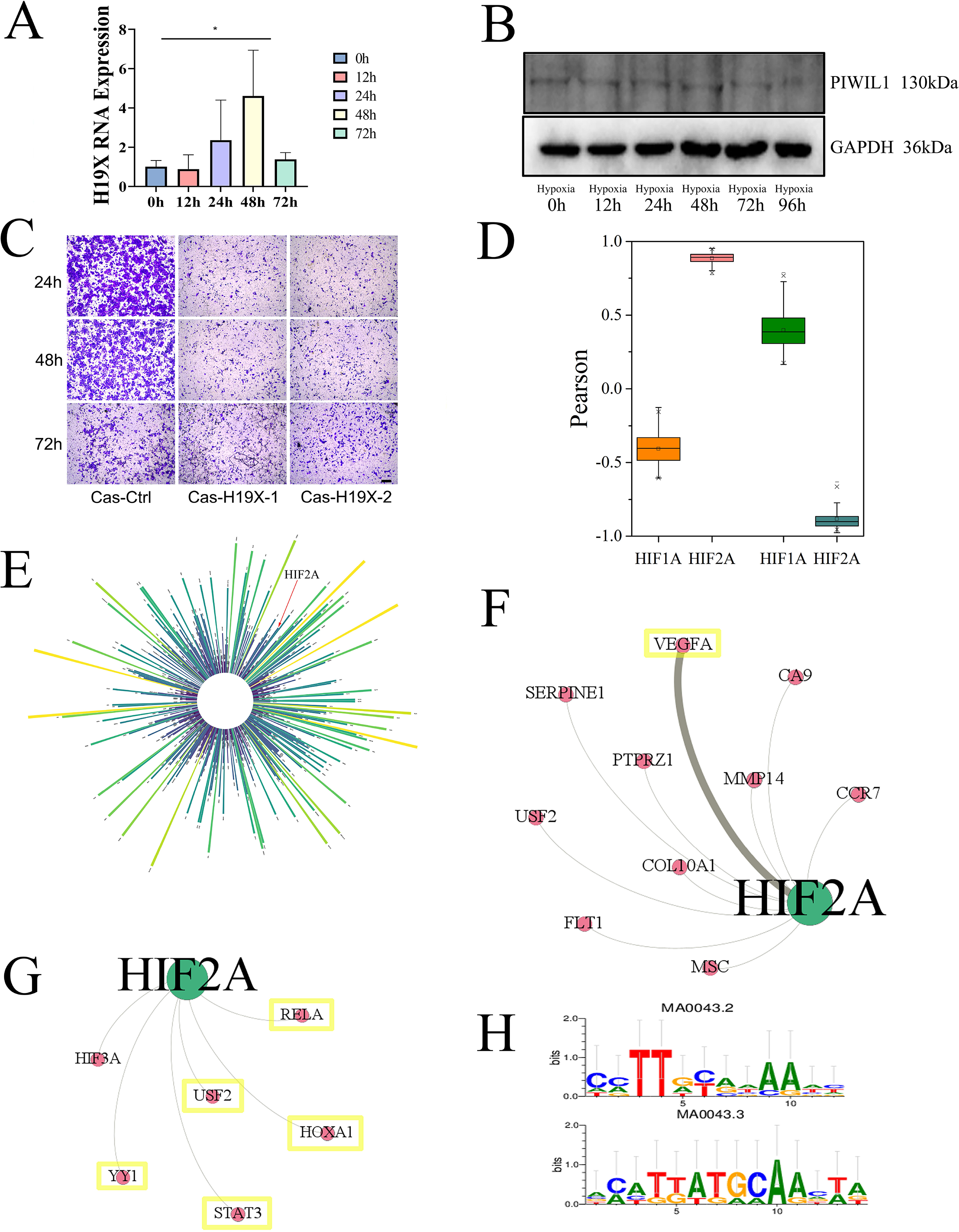
H19X regulates HIF-1A/2A expression, and this reciprocal regulation is amplified under hypoxic conditions. (A) Under hypoxic conditions, H19X expression is significantly upregulated in the first 48 h, and then downregulated after 72 h. (B) Western blot analysis shows that PIWIL1 expression was significantly reduced under hypoxic conditions. (C) A cell invasion assay shows that the number of invading H19X KO cells was significantly reduced under hypoxic conditions. (D) Gene correlation assay shows that HIF-2A is more significantly correlated with H19X targets, including both upregulated and downregulated genes after H19X KO. (E) The promoter sequence of H19X contains multiple transcription factor binding sites, including a HIF-2A site. (F) Network analysis shows that HIF-2A targets the VEGF pathway. (G) The figure shows the transcription factors (TFs) that regulate HIF-2A. The highlighted TFs enable TF binding to the promoter sequence of H19X. (H) The right panel shows the conserved binding motif for HIF-2A, which is found in the H19X promoter region.

Considering co-transcriptional regulation, we used the HTR-8 cell hypoxia treatment model to evaluate the regulation of H19X to HIF-1A/HIF-2A. After hypoxia treatment, HIF-1A/HIF-2A expression increased significantly after 72 h, indicating the successful establishment of a hypoxic model in trophoblasts (Figure 7A, B). We then used the model to evaluate HIF-1A/HIF2A expression following H19X knockout, both at the RNA and protein levels. The results showed that the expression of HIF-1A/HIF2A was significantly reduced in H19X deletion cells (Figure 7C-G). Interestingly, HIF-2A protein expression was more significantly reduced following H19X deletion, indicating that HIF-2A is more closely related to the regulation of H19X (Figure 7D, E). When H19X was deleted in normal conditions and hypoxia, the results showed that HIF-1A was modestly reduced under normal conditions, while under hypoxia, HIF-2A transcripts were more significantly reduced (Figure 7F, G). Taken together, these results show that the regulation of H19X by HIF-2A is stronger after hypoxia, indicating that hypoxia augments H19X regulation by the HIF-2A transcriptional factor, which may have a more significant contribution to the phenotype under *in vivo* conditions. To further investigate whether changes in the expression of HIF impact the VEGF pathway and other downstream pathways, we evaluated the expression of VEGF and downstream genes. The results showed that VEGF transcript and protein levels were significantly downregulated following H19X knockout (Figure 7H, I). We then analyzed transcriptome sequencing data to determine whether the above-mentioned pathways were changed at genomic level. GO analysis indicated that among the downregulated genes, “hypoxia response” and “angiogenesis” were the major pathways in GO terms following H19X deletion (Figure 7J). Analysis of the 20 most significant terms, “viral entry into host cells” and “sprouting angiogenesis” were among the most significant pathways (Figure 7K). Taken together, these results indicate that H19X-HIF-1A/HIF-2A represents a key axis at the transcriptional level, especially under hypoxic conditions.

**Figure 7.**
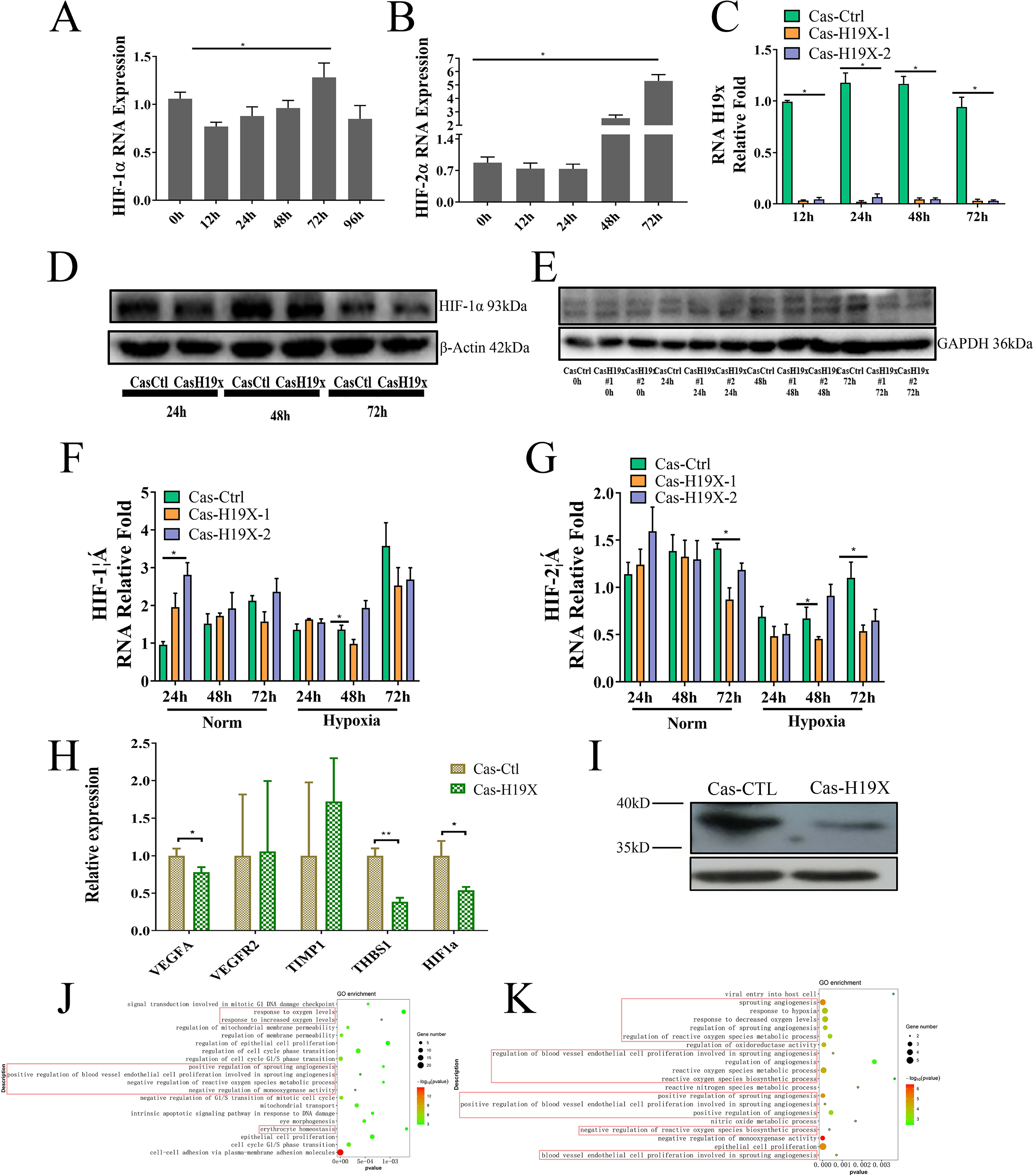
A hypoxia model in trophoblasts confirms that H19X and HIF-2A co-regulation is over-represented in H19X KO cells. (A, B) Real-time PCR shows that hypoxia treatment leads to increased HIF-1A/HIF2A expression. (C) CRISPR/Cas-9 deletion results in diminished expression of H19X under hypoxia. (D, E) Western blotting shows that H19X KO leads to compromised HIF-1A and HIF-2A expression. (F, G) Real-time PCR shows that that HIF-IA and HIF-2A transcript levels are significantly reduced in H19X KO cells. HIF-2A expression is more significantly reduced under hypoxia. *p < 0.05. (H) H19X KO cells also show reduced VEGF mRNA expression. (I) Western blotting shows that VEGF protein is also reduced in the Cas-19X cell line. (J, K) GO analysis of the DEGs shows that hypoxia response and immune activation are the over-represented pathways. Cas-H19X-1 and Cas-H19X-2 are two independent Cas-9 cell lines.

### Knockout of H19X in mice leads to compromised placental development and increased embryo miscarriage under hypoxic conditions

In a previous study, we found that the expression of H19X and PIWIL1 was altered under hypoxic conditions. Hypoxia is an important characteristic of early pregnancy. To investigate whether knockout of H19X affects placental development in the present study, we established a H19X KO mouse model using the CRISPR/Cas-9 method. After genotyping, which showed that H19X-KO mice had a PCR fragment of <1745 bp, one KO line was chosen for functional study. Placental development was evaluated in WT and heterozygous and homozygous H19X knockout mice at E13.5 day. The gross weight of the placenta and the embryo numbers in H19X KO mice were comparable to those in WT mice (Supplementary Figure 14A). Immunohistochemical staining showed that the labyrinth zone length was reduced in KO mice compared to that in WT mice, as was placental development (Figure 8A). The vessel density and branching were significantly reduced, indicating the presence of thrombotic vessels. H&E staining showed a significant reduction in vacuolated foci in the labyrinth of these two groups of mice compared to control mice (Figure 8A). Then, we examined the expression of trophoblast marker genes in the placenta of KO mice on E14.5 day (Gasperowicz *et al*, 2013). Our results showed that expression levels of the Glycogen trophoblast and the spongiotrophoblast marker Tpbpa were significantly reduced, while the expression levels of other markers, including Pcdh12 and Prlgag, showed no significant difference (Figure 8B). We examined the major downstream targets of H19X in the KO mice model and found that the protein expression levels of MIWI and VEGFR2 were significantly reduced (Figure 8C, D). These results indicate that the expression of H19X in extravillous trophoblast (EVT) is important for the interaction of the EVT with the endothelium and is therefore critical for vascular development.

**Figure 8.**
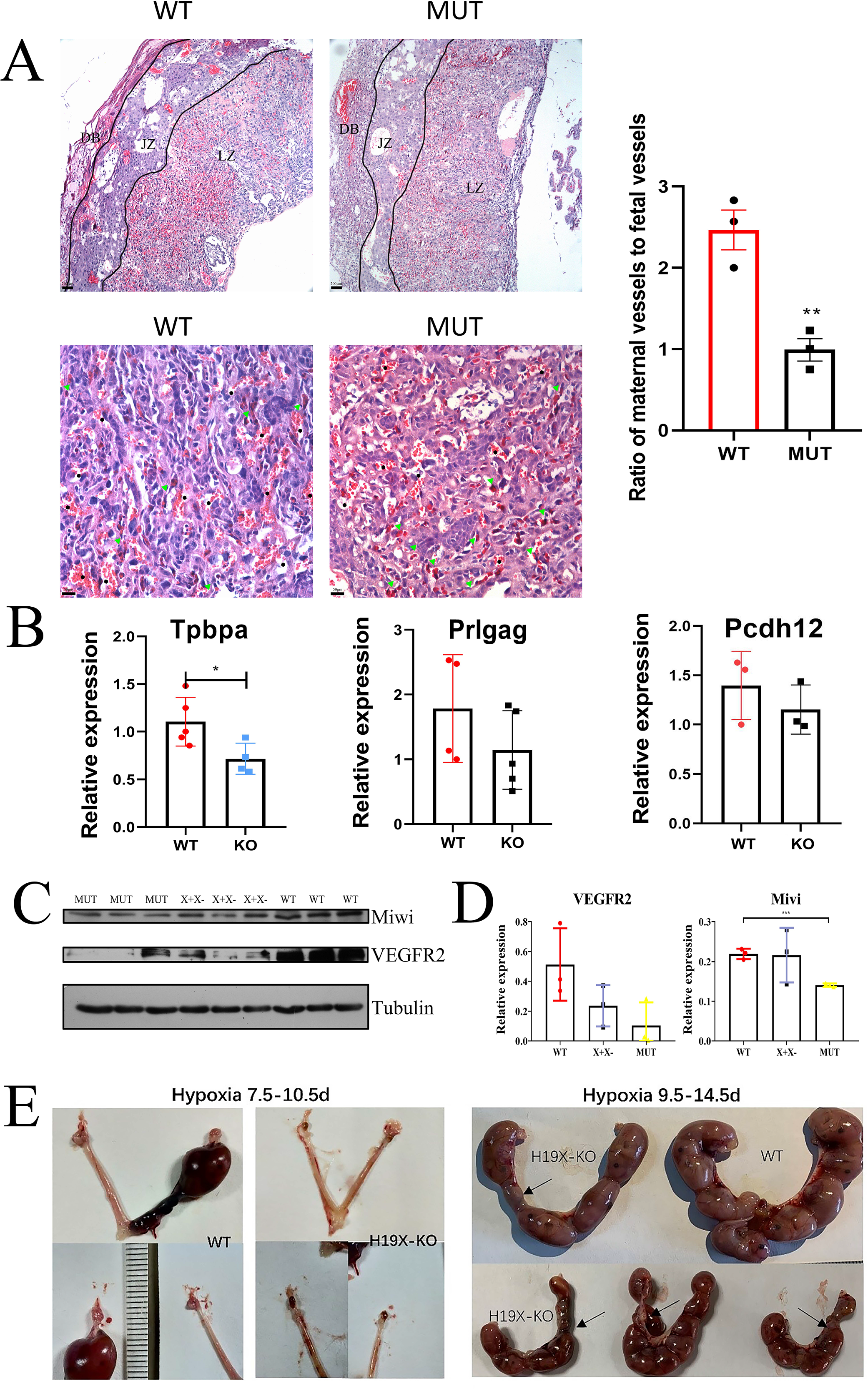
H19X KO mice exhibit poor placental development, angiogenesis defects, and miscarriage under hypoxia treatment. (A) H&E staining of placenta from wild-type (WT) and mutant (MUT) mice at E13.5. Black lines are used to separate the placenta to three sections: the decidua basal (DB), junction zone (JZ), and labyrinth zone (LZ). The LZ is significantly thinner in the MUT placenta than in the WT placenta, as shown in the upper lane. Scale bar, 200 μM. Lower panel: Maternal vessels containing mature red blood cells are labeled with black hexagons, fetal vessels containing mature red blood cells are labeled with **black hexagons**, and fetal vessels containing immature nucleated red blood cells are labeled with green triangles. The ratio of maternal vessels to fetal vessels is significantly decreased in the MUT placenta, as shown in the lower lane. Scale bar, 50 μM. *p < 0.05. (B) Real-time PCR shows that the expression of the Glycongen trophoblast cell marker, Tpbpa, is significantly reduced (p < 0.05), while the expression of other genes, include *Prlgag* and *Pcdh12*, are also reduced, however without significance. (C) Western blotting shows that the expression both MIWI and VEGFR2 is significantly reduced in placenta upon H19X KO. (D) Statistical results. (E) Hypoxia treatment from E7.5 to 10.5 shows abolished embryo implantation compared with the WT control. Hypoxia treatment from E9.5 to 14.5 results in more embryo absorption and miscarriages than the control.

Although placental development appeared to be compromised, the placenta weight and embryo numbers were largely unaffected in the mating experiment. Next, we established a hypoxia model using the KO mice to examine their phenotype under hypoxic conditions. We used two timelines to investigate the effect of 9.5–11.5% hypoxia. The placenta in homozygous mice were completely absorbed, and the size of the ovaries was reduced (Figure 8D). The oxygen concentration varies during different periods of pregnancy. As such, we induced hypoxia from day E9.5 to E14.5 and found that the number of fetuses in the KO mice was reduced compared the number in the control mice. Furthermore, larger number of placenta and fetus were found to be absorbed in the H19X-KO mice, and more variable phenotypes appeared among the fetuses and absorbed fetuses under hypoxic conditions (Figure 8E).

R software was used to analyze the different parameters of placental development, including placenta weight, fetus weight, and the percentage of absorbed fetus, and differences were found between the KO and control mice (Figure 8E). The results showed that the H19X-KO mouse embryos had higher weights. Moreover, the placenta/fetus weight ratio was significantly lower in KO mice than the control, which could be attributed to nutrition exchange between the fetus and the mother, leading to poor development of the placenta and fetus (Supplementary Figure 14). Based on the results of the KO mouse model, high H19X expression may maintain PIWIL1 stability, while the deletion of H19X leads to increased PIWIL1 degradation, affecting the normal development of the placenta. *In vivo* deletion of H19X results in a more significant phenotype under hypoxic conditions.

### H19X-mediated regulation of human trophoblasts is conserved and is relevant to miscarriage

To determine whether H19X plays a role in human primary trophoblast cell invasion, human villi samples were extracted and cell identity was confirmed with the trophoblast markers CK-7 and HLA-G (Figure 9A). Then, H19X was knocked down with both siRNA and shRNA in the primary trophoblast cells. H19X knockdown significantly reduced trophoblast cell invasion (Figure 9B), indicating that H19X plays a role in human primary trophoblast invasion. The RPL sample was then used to determine whether reduced expression of H19X was related to the etiology of repeated pregnancy loss (RPL). Explants of RPL samples showed reduced invasion capacity (Figure 9C). Furthermore, H19X expression was examined in a cohort of RPL samples, and the results showed that H19X expression was significantly reduced compared with that in the control (Figure 9D). Taken together, these results indicate that the function of H19X is conserved between mice and humans. Therefore, H19X dysregulation is associated with the pathogenesis of miscarriage, and the underlying mechanism is shown in Figure 9E.

**Figure 9.**
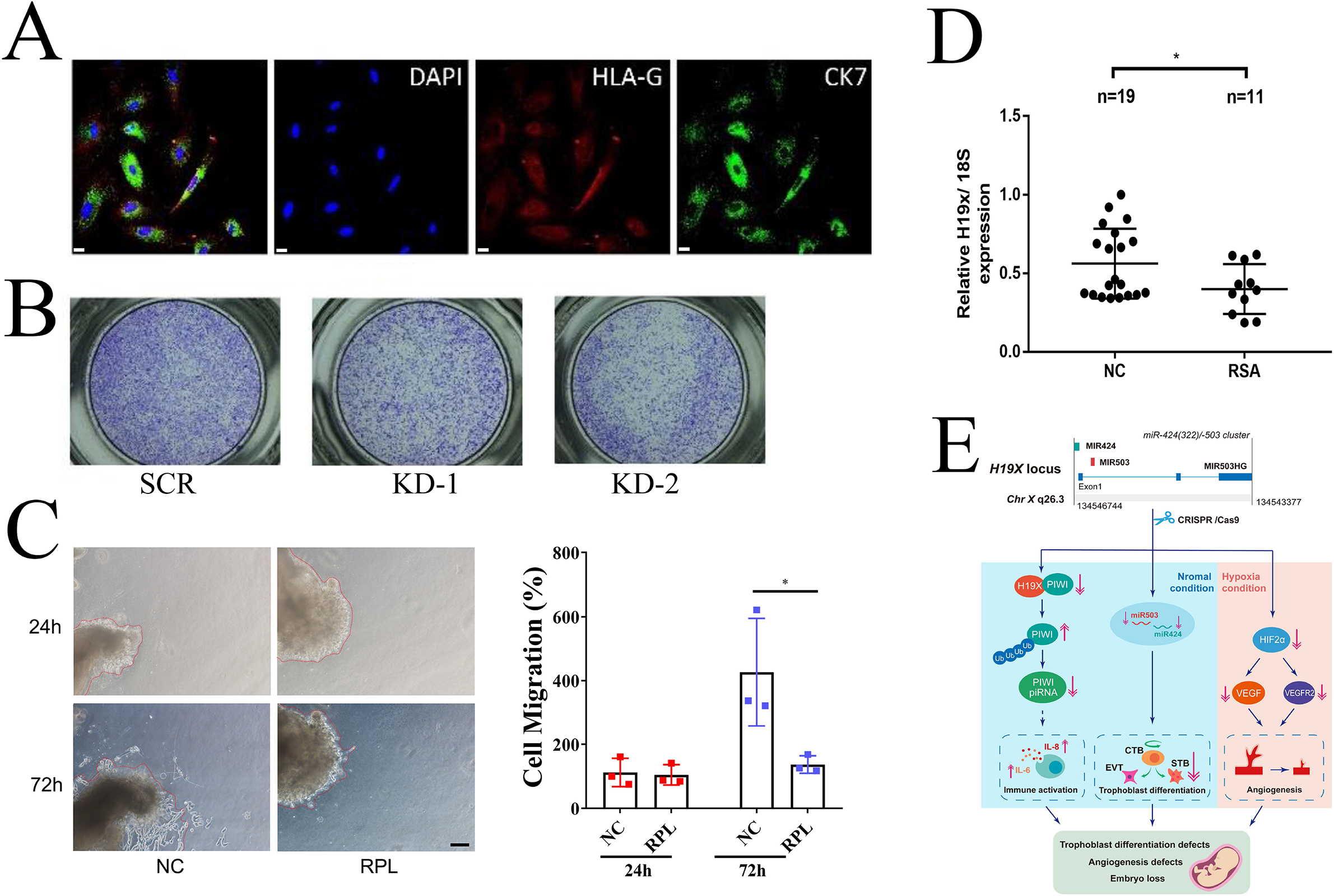
Human samples show that the function of H19X in villi trophoblast cell invasion is conserved. (A) Positive staining of primary trophoblast cells from villi samples for the trophoblast marker CK-7 and HLA-G. (B) Knockdown of H19X in primary trophoblast cells using siRNA. Trophoblast cell invasion is significantly reduced when transfecting siRNA compared with control. (C) Explant of an RPL sample shows reduced invasion capacity compared with the control. Scale bar, 200 μM. Right panel is the statistical results. * means p<0.05. (D) A cohort of RPL samples examined for H19X expression shows that H19X expression is significantly reduced compared with that in the control (p = 0.04) (n = 17, control; n = 11, RPL). (E) Mechanistic insights from the current study. Normal H19X expression is important for PIWIL1 protein stability and miRNA expression, which is important for placental development. Deletion of H19X leads to increased PIWIL1 ubiquitination, which results in piRNA mediated immune activation and dysregulation of miRNA expression, including miRNA-424 and miR-503, which contribute to the development of placental defects. Under hypoxia, the feedback loop involving H19X and HIF-2A is amplified, leading to more significantly reduced expression of HIF-2A, compromised expression of VEGF and VEGFR2, and angiogenesis defects. This phenotype became significant under hypoxia.

## Discussion

In the current study, H19X was found to play a critical role in placental development by coordinating miRNA and piRNA expression. We also found that a feedforward loop between H19X and HIF-2A was amplified under hypoxic conditions, resulting in poor placental development. Notably, H19X was found to interact with PIWIL1 in trophoblasts, and may thus play a critical role in trophoblast cell invasion. This study is the first to examine the expression of PIWIL1 in the placenta, which plays a role in trophoblast behavior. PIWIL1 is a well-known germ cell-expressed protein whose major functions include repressing retrotransposons and regulating target gene expression and recent study showed that PIWIL1 is involved in tumorigenesis via a piRNA-independent mechanism (Li *et al*, 2020). Interestingly, A recent bioinformatic study showed that one group of piRNAs are potentially implicated in functional regulation of the placenta (Chirn *et al*, 2015). The present study provides the first experimental evidence that PIWIL1 is expressed in the placenta and plays important roles in trophoblast immune responses by regulating piRNA expression. Through a bioinformatic analysis, we found that deletion of H19X resulted in PIWIL1 degradation, leading to piRNA dysregulation, which, in turn, is related to immune activation. In accordance with previous studies, our findings show that the key functions of piRNAs are immune surveillance and immune suppression (Robinson *et al.*, 2020; Wang *et al*, 2020). In addition, a recent study showed that some germ cell-specific genes such as *TEX19.1* also play important roles in placental development (Judith *et al*, 2013). It is also possible that PIWIL1 represses retrotransposons during placental development (Teixeira *et al*, 2017).

In the present study, H19X was found to regulate the stability of PIWIL1 by inhibiting the ubiquitination of PIWIL1. During spermiogenesis, PIWIL in germ cells is degraded through APC-mediated ubiquitination (Zhao *et al.*, 2013). This study is the first to show that the lncRNA H19X is involved in inhibiting the ubiquitination of PIWIL1. Further study is needed to determine whether degradation of PIWIL induced by ubiquitination of PIWIL1 requires APC. MIWI female mice are known to have normal fertility and a normal number of pups. However, it should be noted that there are multiple MIWI homologues in mice. Therefore, *in vivo* compensation of other MIWI homologue in MIWI mice during placental development remains possible. Nevertheless, it remains to be shown whether MIWI mice have compromised placental development, at least under hypoxic conditions.

The relationship between the lncRNA H19X and its host miRNAs, miR-424 and miR-503, represents another level of regulation. Although a recent *in vitro* study showed that H19X plays a role in trophoblast migration (Wang *et al.*, 2019b), the author only showed that miR-503HG (a transcript of H19X) played a role in trophoblast migration, but no mechanistic insight was gleaned from the study. In the present study, using the trophoblast cell line HTR-8, we showed that deletion of H19X also compromised the expression of several miRNAs, including that of miR-503 and miR-424, and reciprocal regulation between miR-424 and H19X via transcription. Thus, H19X and miR-424 represent another example of reciprocal regulation involving miRNAs and lncRNAs in trophoblast differentiation.

Both our *in vitro* and *in vivo* data show that H19X knockout results in the dysregulation of trophoblast behavior when compared with the control. However, this phenotype becomes more marked under hypoxic conditions. The transcriptome results showed that H19X deletion leads to hypoxia-related changes in the transcriptome, and the experimental data support the observation that under hypoxic conditions, the deletion of H19X leads to downregulation of HIF-1A and HIF-2A, which in turn leads to defects in angiogenesis, with compromised VEGF and VEGFR2 expression. HIF-2a is a critical transcription factor for embryo implantation in mice (Highet *et al*, 2015; Matsumoto *et al*, 2018). The mechanism underlying the regulation of HIF1A/HIF2A by H19X is likely to involve co-transcriptional regulation and warrants further investigation (Wang *et al*, 2019a). Since H19X is expressed in both cytoplasm and nucleus, and our bioinformatic analysis showed that both H19X and HIF-2A share co-transcription factors, it is likely that H19X is regulated through a cis-acting mechanism at the HIF-2A promoter. Under hypoxic conditions, feedforward regulation is amplified, which leads to decompensation of HIF-2A. As a result, diminished expression of VEGF and VEGFR2 leads to angiogenesis defects. The major mechanism by which H19X modulates HIF-2A expression is shown in Figure 9E.

The main function of the placenta is to regulate offspring development. It has been shown that lncRNAs play a critical role in offspring development. In particular, as one of H19X related lncRNA, lncRNA H19 has been shown to be involved in placental development, by controlling placental growth via miR-675 (Keniry *et al.*, 2012). Deletion of H19 and miR-675 results in excessive placental and fetal growth. In our *in vivo* model, deletion of H19X resulted in excessive fetal growth and fetal loss. The corresponding *in vivo* data suggest that H19X plays a critical role in the exchange of nutrition between mother and offspring across the placenta. Therefore, the *in vivo* function of H19X is similar to that of H19 in the control of trophoblast growth as well as nutrition exchange in the LZ region. However, the exact mechanism by which H19X regulates fetal growth warrants further investigation.

Poor placentation is observed in various pregnancy-related diseases, including preeclampsia, miscarriage (Burke *et al*, 2016), and recurrent pregancy loss (RPL. These devastating diseases affect 3–6% of the pregnant women worldwide (Tang *et al*, 2017; Troy *et al.*, 2012). Increased inflammation and/or angiogenesis defects constitute the major pathogenetic pathways of poor placentation-related diseases (Kumar & Goyal, 2017). This study is the first to show that a placenta-specific lncRNA may play a critical role in placental development. Blocking the production of H19X was found to result in an imbalance in nutrient exchange across the placenta in a mouse model. lncRNAs are being extensively studied owing to their vast potential as drug targets for immunological diseases, cardiovascular diseases, and cancers (Atianand & Fitzgerald, 2016). Since H19X is a placenta-specific lncRNA that plays critical roles in trophoblasts, it could be a viable target for placenta-related disease (Pachera *et al*, 2020; Yu *et al*, 2018). Elucidating the mechanism underlying the actions of H19X and PIWIL1 could pave the way for the development of treatments for placenta-related disease in the near future.

## Materials and Methods

For an expanded version of the Materials and Methods section, please refer to the online-only data supplement.

### CRISPR/Cas-9 deletion of H19X in mouse model

A Gm28730 knockout in C57BL/6 mice was established using CRISPR/Cas-mediated genome engineering. The mouse Gm28730 gene (Ensembl: ENSMUSG00000101603; Havana transcript: OTTMUST00000121000) is located on the X chromosome. Exon 2 was selected as the target site in mouse Gm28730. Cas9 mRNA and gRNA generated by *in vitro* transcription was co-injected into fertilized eggs to establish KO mice. The founder mice were genotyped by PCR and DNA sequencing analysis. The positive founders were bred to the next generation, which was genotyped by PCR and DNA sequencing analysis. All chemicals and reagents were of analytical grade. C57BL/6 mice were purchased from Chengdu Dashuo Biological Institute (Chengdu, China).

### Cell culture

Human extravillous trophoblast (HTR-8/SVneo) cell lines were cultured in RPMI-1640 medium at 37°C in a 5% CO_2_ humidified environment (Thermo Fisher Scientific, USA). The media contained 10% FBS, 100 U/mL penicillin, and 100 mg/mL streptomycin.

### Cell proliferation and invasion assay

Cells were seeded into 24-well plates at a concentration of 20,000/well on day 0. Then, at 24, 48, 72, and 96 h after seeding, four wells of the cells were digested with trypsin and counted using a cell counting chamber. The cell proliferation curve was created using Prism software.

Cell invasion assays and *in vitro* angiogenesis assays were performed as described previously (Yang *et al.*, 2016).

### Plasmids structure

Full-length human H19X (806 bp) was amplified by PCR using Fast *Pfu* DNA polymerase (TransGen Biotech, Beijing, China) from HTR-8/SVneo cDNA before linking to the pcDNA3.1 vector. PIWIL1 cDNA (gifted by Dr. Qintong Li) was subcloned into pcDNA3.1. Part of human H19X Exon3 was cloned into pcDNA3.1 to produce templates for *in vitro* transcription. All primers used are listed in Supplementary Table 1.

### RNA immunoprecipitation (RIP)

RNA immunoprecipitation was performed using the Magna RIP RNA-Binding Protein Immunoprecipitation Kit (17-700; Millipore) according to the manufacturer’s instructions. Briefly, 1 × 10^7^ HTR-8/SVneo cells were homogenized using RIP lysis buffer plus RNase inhibitor (Thermo Fisher Scientific) and a proteinase inhibitor cocktail (Roche). After soft-freeze thawing, the mixtures were incubated with anti-PIWIL1 antibody (5 μg; ab181056; Abcam) or isotype IgG (5 μg; ab172730; Abcam), and Protein A/G beads overnight at 4°C. After washing six times with RIP wash buffer, the RNA-protein complex-associated beads were incubated in proteinase K buffer. The RNA was isolated by salt precipitation according to the manufacturer’s instructions, followed by qRT-PCR analysis.

### RNA pull-down assay

An RNA pull-down assay was performed as previously described, with some modifications (Feng *et al*, 2014). Briefly, a biotin-labeled H19X RNA molecule and its antisense RNA molecule (X91H) were transcribed *in vitro* from linear plasmid templates by T7 RNA polymerase (NEB, UK) with Biotin pre-labeled Uridine (Roche) and other reagents according to protocol for T7 RNA polymerase. The corresponding products were then purified using an RNA Extraction Kit (Omega). For each group, 1 × 10^7^ HTR-8/SVneo cells were harvested using a regular trypsinization technique, followed by homogenization using a Dounce homogenizer, before washing twice with PBS. Briefly, 50 μL of streptavidin magnet beads was pre-washed three times in NT2 buffer, and then resuspended in NT2 buffer. A sufficient amount of pre-washed beads were then added to lysed HTR-8/SVneo cells to pre-clear the whole cell lysate before separating into a magnetic frame. The supernatant was then transferred to a new tube. The protein concentration in the pre-cleared lysate was determined by BCA protein assay (Thermo), and 4% of the lysate was saved as input. Biotinylated RNA (10 pmol) was diluted into the RNA structure buffer, heated to 90°C for 2 min, and immediately transferred to ice for 2 min, then incubated at room temperature to allow the formation of RNA secondary structure. The resulting structured RNA was added to 200 μg of pre-cleared lysate supplemented with 0.1 μg/μl tRNA and incubated for 2 h at 4°C. Then, 50 μL of pre-washed streptavidin magnetic beads were added, and the mixture was incubated for another 2 h at 4°C. After incubation, the protein-RNA-beads complexes were washed five times with NT2 buffer, and then mixed with 2× Laemmli loading buffer. After boiling for 10 min, the supernatant was loaded and separated by 8% SDS PAGE, and then stained with Coomassie Brilliant Blue to detect H19X-specific binding proteins. Specific bands were cut out of the gel for analysis by mass spectrometry. The NT2 buffer and RNA structure buffer were prepared as described previously(Feng *et al.*, 2014).

### Immunofluorescence (IF)

HTR-8 cells were fixed with 4% PFA and treated with 0.3% Triton-X100. After washing the slides with PBS, they were incubated overnight at 4°C with primary antibodies against PIWIL1 (ab181056; Abcam). The cells were then washed with PBS and incubated with Alexa Fluor secondary antibodies (Life Technologies) for 60 min. The slides were stained with DAPI (1 μg/mL) for 5 min after washing with PBS three times. The results were analyzed using the FV1000 Confocal system (Olympus, USA).

### Immunohistochemical (IHC), cytokine array and ELISA

#### Human cytokine antibody array

The levels of various immune factors in condtinal medium (CM) were detected using the Human Kit (R&D) according to the manufacturer’s instructions. Briefly, 200 μl aliquots of samples were mixed with reconstituted detection antibody cocktail. After incubation, the membranes were treated with 1.5 mL of the sample/antibody mixture overnight at 4°C on a rocking platform shaker, followed by three washes. Then, the membrane was incubated with streptavidin-horseradish peroxidase solution, and washed three additional times. Finally, signals were generated by exposure to the ECL chemiluminescent substrate supplied in the kit.

IHC staining was performed on 5 μm-thick paraffin-embedded tissue sections. The slides were dried overnight and then immersed in xylene for 20 min. Next, the sections were dehydrated in graded ethanol (100%, 95%, 90%, 80%, 75%, and 50%) for 5 min, followed by the steps described in the manufacturer’s protocol (Zhongshanjinqiao Co., Beijing, China).

### *In situ* hybridization (ISH) and fluorescence *in situ* hybridization (FISH)

A digoxin-labeled human H19X probe was transcribed *in situ* from a linear template plasmid by T7 RNA polymerase (NEB, UK) with Digoxin-labeled Uridine (Roche) and other reagents according to the instructions for T7 RNA polymerase, and then purified using the blood RNA extraction kit (Omega).

For the FISH and ISH staining of lncRNA, cryosections of trophoblasts and HTR-8 cells at 50–70% confluence were washed with DEPC-PBS, fixed in 4% PFA, and subsequently immersed in 0.1 mol/L triethanolamine containing 0.25% acetic anhydride for 10 min at 37°C. The cells were then permeabilized by incubation with 0.2 mol/L HCl for 10 min after washing with PBS, and treated with proteinase K (1 mg/mL) at 37°C for 10 min. The cells were then incubated in pre-hybridization buffer, followed by hybridization with the probe at 60°C in a humid oven covered with a lid overnight. During pre-hybridization, 1 ng/mL of probe was added, denatured by heating at 85°C for 10 min, and then immediately cooled on ice for at least 5 min. The following day, the slides were washed in 0.1× SCC containing 50% formamide, and then incubated at 60°C for 30 min twice. Then, the slides were washed with 2× SSC at 37°C. After blocking with blocking buffer, the slides were immersed in anti-Digoxin-Rhodamine Fab fragments (1 mg/mL; Roche) antibody or anti-Digoxin-AP (Roche) for 1 h or 1 day at 37°C, respectively. After counterstaining the slides with DAPI (1 mg/mL) or nuclear fast red, the slides were covered with Prolong Gold antifade reagent (1620476; Life Technologies, USA). The resulting signal was detected using a confocal microscope (FV1000; Olympus, Japan).

### CRISPR/Cas9 in cultured cells

CRISPR/Cas9 was used to knockout the third exon of H19X in HTR-8/SVneo cells, as previously described (14). Briefly, sgRNA sequences targeting the flank of H19X Exon3 were obtained from http://crispr.mit.edu/. The sgRNA-Cas9 plasmids were constructed using the PX459 vector. After co-transfection with the two sgRNA-Cas9 plasmids containing sequences flanking H19X Exon3 for 48 h, the cells were incubated with 1 μg/ml puromycin for 2 days to select successfully transfected cells. The cells were then digested with trypsin. After measuring their concentration, the cells were diluted to a single cell per 100 μL of medium and re-seeded onto 96-well plates. The single cell-derived colonies were observed after re-seeding cells for 1 or 2 weeks. Once the colonies had grown enough, some were selected for genotype examination. H19X E3 knockout cell colonies were continually cultured for follow-up experiments. Untreated HTR-8/SVneo cells from the same passage were used as a knockout control, denoted as Cas-CTL. The sequences of oligos and PCR primers used are listed in **Supplementary Table 1**.

### Co-immunoprecipitation (Co-IP)

Cas-H19X and Cas-CTL HTR-8/SVneo cells were transfected with 4× Flag-Ubi-pcDNA3.1 for 24 h, treated with the proteasome inhibitor MG132 (at 20 mmol/L) for 4 h, and lysed using RIPA plus proteinase inhibitor cocktail (Roche). Each lysate, containing 150 μg protein, was added to 30 μL of pre-washed protein A magnet beads (Thermo Fisher Scientific) and 1 μg of PIWIL1 antibody (ab181056; Abcam), incubated overnight, and then separated using a magnetic frame. After sufficient washing, the protein-bead complexes were mixed with SDS loading buffer and then subjected to an ordinary western blot assay. PIWIL1 antibody (ab181056; Abcam) and FLAG antibody (Zen-Bio, Chengdu, China) were used to detect the proteins.

### Western blot analysis

The expression levels of PIWIL1, HIF-1A, HIF-2A, and VEGF-Ain HTR-8/SVneo cellswere measured by western blotting. HTR-8/SVneo cells were separately seeded onto 6-well plates at a density of 5 × 10^5^ cells/well and were homogenized with cell lysis buffer containing protease inhibitors. Then, total protein samples from different cells were separated by 10% SDS-PAGE. The proteins in the gels were transferred to polyvinylidene difluoride membranes and incubated with rabbit anti-mouse GAPDH or. After 24 h, the membranes were incubated with anti-rabbit antibodies and detected with Immobilon Western HRP Substrate (Millipore, USA) on a Bio-Rad ChemiDoc MP System (Bio-Rad Laboratories, USA).

### Animal care and mating

All animal experiments were performed according to the guidelines of the Animal Experimentation Ethics Committee of Sichuan University. To establish pregnant mice, 8–12-week-old female C57 BL/6 mice were mated overnight with males. The presence of a vaginal plug was defined as 0.5 d of gestation. Pregnant females were administered liposomes or free siRNA at a dose of 0.35 mg/kg via tail vein injection.

### Animal experimental protocol

To analyze pregnancy outcomes, mice were euthanized at 18.5 d of gestation. The number of live pups per mouse, placental weight, and fetal weight were recorded. For histological analysis, placentas and kidneys were collected. Placental histology was performed after hematoxylin and eosin (H&E), periodic acid-Schiff (PAS), and VEGF staining. Kidney histology was performed after H&E staining.

### Statistical analysis

All data are expressed as the mean ± standard error of the mean (SEM). All statistical analyses were performed using GraphPad Prism software (version 5.0; GraphPad, USA). For two-group comparisons, a two-tailed Student’s t-test was used. Statistical comparisons of more than two groups were performed by one-way analysis of variance (ANOVA) for multiple groups. Significant differences between or among groups are indicated by *p < 0.05, **p < 0.01, and ***p <0.001, respectively.

### RNA isolation, library preparation, and sequencing

#### Small RNA sequencing

Total RNA (3 μg) from the placenta of each sample was used as the input material for the small RNA library. RNA degradation and contamination were monitored by separation on 1% agarose gels. RNA purity was verified using a NanoPhotometer^®^ spectrophotometer (IMPLEN, USA). RNA was quantified using the Qubit^®^ RNA Assay Kit and a Qubit^®^ 2.0 Fluorometer (Life Technologies, USA). RNA integrity was assessed using the RNA Nano 6000 Assay Kit. Sequencing libraries were generated using the NEBNext^®^ Multiplex Small RNA Library Prep Set for Illumina^®^ (NEB, USA) according to the manufacturer’s instructions. Index codes were added to attribute sequences to each sample. Clustering of the index-coded samples was assessed on a cBot Cluster Generation System using the TruSeq SR Cluster Kit v3-cBot-HS (Illumina) according to the manufacturer’s instructions. After cluster generation, the library preparations were sequenced on an Illumina HiSeq 2500/2000 platform, and 50-bp single-end reads were generated.

#### RNA-sequencing analysis

RNA sequencing (RNA-seq) analysis was performed using the total RNA purified from adult placenta using TRIzol reagent. Libraries were multiplexed using NEBNext^®^ Multiplex Oligos for Illumina^®^, and reads were sequenced on an Illumina HiSeq 3000.

### Data analysis

#### Small RNA analysis

The small RNA-seq data were uploaded to the GEO database (GSE158462). All reads were trimmed before aligning to the genome (Supplementary Figure 1). We used Trim Galore (https://github.com/FelixKrueger/TrimGalore/releases) to filter out low quality reads and reads that contained adapters. The small RNA-seq was mapped to the reference sequence using Bowtie (B & M, 2009), without mismatches, to analyze their expression and distribution on the reference. The files were converted to the bam format in SAMtools 1.7 (Li *et al*, 2009). We downloaded the coordinates and annotations of the miRNAs from the UCSC Genome Browser (https://genome.ucsc.edu/). The genome files and annotation files for piRNAs were downloaded from piRNAdb (https://www.pirnadb.org/download), and duplicates were removed using Picard tools (http://broadinstitute.github.io/picard/). Reads mapped to regions of each miRNA/piRNA were counted using featureCounts (Liao *et al*, 2014), and then the reads per kilobase per million fragments (RPKM) was calculated. Genes that were differentially expressed between the experimental and control groups were identified using the edgeR package (Robinson *et al*, 2010). The differentially expressed miRNAs/piRNAs were selected using a threshold of P < 0.05 and a fold change ⩾1.

MiRwalk (Sticht *et al*, 2018) was used to predict the target genes of the differentially expressed miRNAs. Here, the minimum number of nucleotides in an miRNA sequence through which they can make one or more interactions with possible targets was set to seven (P < 0.05), and predictions were made by comparative analysis using four programs (miRanda, miRDB, miRWalk, and TargetScan). For differentially expressed piRNAs, miRanda (v3.3a) (Enright *et al*, 2003) was used to predict target genes. There was overlap between the target genes of differentially expressed piRNAs and miRNAs. Gene Ontology (GO) enrichment analysis was performed on the predicted target genes of the differentially expressed miRNAs/piRNAs. The overlapping genes with piRNAs were extracted using bedtools (Quinlan, 2014), and a functional enrichment analysis was performed for these genes. The gene set enrichment analysis was performed using clusterProfiler in the R package (Yu *et al*, 2012) and Metascape (http://metascape.org) (Zhou *et al*, 2019). REViGO (Supek *et al*, 2011) was used to remove redundant GO terms and summarize them.

The obtained Bam file was used to visualize the high-throughput sequencing data at H9X, miR424, and miR503 in the IGV browser.

#### RNA-seq analysis

The raw data were filtered with Trim Galore to obtain high-quality (clean) data. FastQC was used to evaluate data quality. Clean reads were aligned to the human genome (hg38) by TopHat (Trapnell *et al*, 2009) with the parameter “--read-mismatches=2 --read-gap-length=2.” The mapping files were converted to the bam format available in SAMtools 1.7 (Li *et al.*, 2009). Differentially expressed genes between groups were analyzed by DESeq (Anders *et al*, 2010). Using the two criteria of a |log2(fold change)| >1 and padj <0.05, the differentially expressed genes between samples were identified. Gene ontology (GO) analyses of differentially expressed genes were performed using clusterProfiler (Yu *et al.*, 2012), using a P-value cutoff of 0.05 for statistically significant enrichment.

### HIF1A/HIF2A analysis

To identify correlations between HIF1A, HIF2A, and DEGs, Pearson’s correlation coefficient was used. The correlation between DEGs and HIF2A was more obvious. Therefore, we focused on HIF2A and examined target genes regulated by HIF2A in TRRUST (version 2) (Heonjong Han 1 2018). In TRRUST, some TFs that regulate HIF2A have been identified. JASPAR was used to identify transcription factor-binding sites in the H19X promoter sequence (Fornes *et al*, 2020). The predicted sequence of HIF2A was extracted, and sequence logos were drawn using WebLogo (Crooks *et al*, 2004).

## Acknowledgements

We would like to thank the Prof.Fu Xiang hui to constructive comments. This work was supported by the National Key R&D research program (2018YFC1002804) and the National Natural Science Fund of China (81771642)

## Author contributions

L. T.T contributed the bioinformatic analysis; Y.Y contributed the hypoxia treatment work; T.L, W.K and Y.M contributed the cell culture; L.J, T.L, and H.G.L contributed the molecular biology work and immunohistochemistry; C.Y.C, Z. Y.T and H.J.B contributed the animal model; X. W.M designed the experiment and analyzed the data, wrote the manuscript; Z.X.M wrote the manuscript.

## Data Availability Section

Data availability

The datasets produced in this study are available in the following databases: RNA-Seq data: Gene Expression Omnibus GSE158462 (https://www.ncbi.nlm.nih.gov/geo/query/acc.cgi?acc=GSE158462).

All relevant data that support the findings of this study are available from the correspondence author upon reasonable request.

